# Werner syndrome protein (WRN) regulates cell proliferation and the human papillomavirus 16 life cycle during epithelial differentiation

**DOI:** 10.1101/2020.05.18.103309

**Authors:** Claire D. James, Dipon Das, Ethan L. Morgan, Raymonde Otoa, Andrew Macdonald, Iain M. Morgan

**Author notes:** Tumor Biology Section, National Institute on Deafness and Other Communication Disorders, NIH, Bethesda, MD 20892, USA. These authors contributed equally to the work (Claire D. James, Dipon Das).

## Abstract

Human papillomaviruses recruit a host of DNA damage response factors to their viral genome to facilitate homologous recombination replication in association with the viral replication factors E1 and E2. We previously demonstrated that SIRT1 deacetylation of WRN promotes recruitment of WRN to E1-E2 replicating DNA, and that WRN regulates both the levels and fidelity of E1-E2 replication. The deacetylation of WRN by SIRT1 results in an active protein able to complex with replicating DNA, but a protein that is less stable. Here we demonstrate an inverse correlation between SIRT1 and WRN in CIN cervical lesions when compared with normal control tissue, supporting our model of SIRT1 deacetylation destabilizing WRN protein. We CRISPR/Cas9 edited N/Tert-1 and N/Tert-1+HPV16 cells to knock out WRN protein expression and subjected the cells to organotypic raft cultures. In N/Tert-1 cells without WRN expression there was enhanced basal cell proliferation, DNA damage and thickening of the differentiated epithelium. In N/Tert-1+HPV16 cells, there was enhanced basal cell proliferation, increased DNA damage throughout the epithelium and increased viral DNA replication. Overall, the results demonstrate that the expression of WRN is required to control the proliferation of N/Tert-1 cells and controls the HPV16 life cycle in these cells. This complements our previous data demonstrating that WRN controls the levels and fidelity of HPV16 E1-E2 DNA replication. The results describe a new role for WRN, a tumor suppressor, in controlling keratinocyte differentiation and the HPV16 life cycle.

**Importance:** HPV16 is the major human viral carcinogen, responsible for around 3-4% of all cancers worldwide. Our understanding of how the viral replication machinery interacts with host factors to control/activate the DNA damage response to promote the viral life cycle remains incomplete. Recently, we demonstrated a SIRT1-WRN axis that controls HPV16 replication and here we demonstrate that this axis persists in clinical cervical lesions induced by HPV16. Here we describe the effects of WRN depletion on cellular differentiation with and without HPV16; WRN depletion results in enhanced proliferation and DNA damage irrespective of HPV16 status. Also, WRN is a restriction factor for the viral life cycle as replication is disrupted in the absence of WRN. Future studies will focus on enhancing our understanding of how WRN regulates viral replication. Our goal is to ultimately identify cellular factors essential for HPV16 replication that can be targeted for therapeutic gain.

## Introduction

HPV are causative agents in around 5% of all cancers, including over 90% of cervical and 70-80% of oropharyngeal (1). Following infection of basal epithelial cells, cellular factors binding to the Long Control Region (LCR) induce viral transcript expression (2). This transcript is then processed and translated to produce the viral proteins. The major viral oncogenes E6 and E7 target, among other host proteins, p53 and pRb function respectively, resulting in enhanced proliferation of the infected cell (3). Two viral proteins are required for replication of the viral genome: E1 and E2 (4). E2 is a DNA binding protein that forms homo-dimers and binds to 12bp palindromic sequences in the LCR surrounding the A/T rich origin of viral replication (5). Via a protein-protein interaction, E2 recruits E1 to the origin of replication and E1 then forms a di-hexameric complex and interacts with host DNA polymerases to regulate replication of the viral genome (6). Following initiation of replication the viral genome establishes in the infected cell at 20-50 copies per cell. The proliferating cell then migrates through the epithelium, maintaining 20-50 genome copies per cell. In the upper layers of the epithelium, the viral genome undergoes amplification to around 1000 copies per cell and at this stage, the structural L1 and L2 proteins are expressed. L1 and L2 then encapsulate the viral genome forming infectious particles that egress from the upper layers of the epithelium (7-9).

During the initial establishment phase of viral replication there is torsional stress on the 8kbp genome as it produces the 20-50 copies per cell that establish infection (10). This creates the formation of damaged DNA structures on the replicating genome and activation of the DNA damage response (DDR). HPV proteins E7 and E1 can also directly activate the DDR when overexpressed in epithelial cells (11-20). The HPV cells proliferate with an active DDR, an unusual outcome as activation of the DDR by external agents ordinarily promotes a cell cycle arrest in order to repair host DNA damage mediated by the external agent (21). This active DDR in the HPV infected cells is required for the amplification stage of the viral life cycle; disruption of the ATM and ATR DDR kinases blocks the amplification stage (21). The replicating viral DNA recruits a number of DDR proteins to promote replication of the viral genome, a process proposed to occur via homologous recombination (HR). The MRN (MRE11-RAD50-NBS1) complex is among the first proteins recruited to damaged DNA, and all three components of MRN are recruited to the viral genome (22, 23). Additional factors involved in HR are recruited to the viral genome, including RAD51 and BRCA1 (24). Many of these DDR proteins are essential for completion of the HPV life cycle.

Our lab has focused on identifying cellular proteins that interact with the viral replication factors E1 and E2 as a means of enhancing our incomplete understanding of host-pathogen interactions that are essential for HPV life cycles, using HPV16 as a model. HPV16 is responsible for around 50% of HPV positive cervical cancers and 80-90% of HPV positive oropharyngeal cancers. We have identified several proteins as being involved in E1-E2 DNA replication including TopBP1, SIRT1 and WRN, all proteins that have key roles in the DNA damage response, HR and DNA replication (25-30). Our recent work on WRN demonstrated that deacetylation of this protein by SIRT1 (a protein that also deacetylates TopBP1) promotes binding of WRN to HPV16 origin containing plasmid DNA being replicated by E1-E2 (25). CRISPR/Cas9 removal of SIRT1 expression results in hyper-acetylation and stabilization of WRN. However, this acetylated WRN is unable to efficiently complex with E1-E2 replicating DNA, as determined by chromatin immunoprecipitation. WRN has 3’ to 5’ exonuclease and helicase activity and is involved in multiple DNA damage and repair processes including non-homologous end joining (31-34). However, it is also involved in HR and replication, and as this is the process used by E1-E2 to replicate the viral genome, the role of WRN in viral replication is to promote HR (32, 35-43). In the absence of SIRT1, E1-E2 replication has a reduced fidelity and one reason for this could be the lack of WRN recruitment to the replicating DNA (26). CRISPR/Cas9 editing of WRN from C33a cells resulted in a similar phenotype to that of SIRT1 deletion: increased DNA replication that has a reduced fidelity (25). Therefore, a SIRT1-WRN axis controls recruitment of WRN to E1-E2 replicating DNA, and a lack of SIRT1 or WRN results in mutagenic HPV16 E1-E2 DNA replication.

The E1-E2 replication assays were carried out in C33a cells, a HPV negative cervical cancer cell line routinely used for viral replication and transcription assays. In this report we expand our investigation of WRN in the HPV16 life cycle. We demonstrate that in CIN1, 2 and 3 lesions there is an increasing tendency for elevated SIRT1 expression and reduced WRN expression. This inverse correlation of expression of these proteins reflects what we observe in our C33a studies where SIRT1 activity deacetylates and destabilizes WRN, although the deacetylated WRN is active in interacting with the E1-E2 replicating DNA and promoting high fidelity DNA replication (25). Others have also demonstrated regulation of WRN function by SIRT1 deacetylation (44-46) SIRT1 is essential for the HPV31 life cycle in cervical keratinocytes, demonstrated by using shRNA to eliminate SIRT1 expression (30). We therefore moved on to investigate the role of WRN during the HPV16 life cycle. We have established a model system in our lab for the HPV16 life cycle using N/Tert-1 cells (47-49). In this system, N/Tert-1+HPV16 cells are transcriptionally reprogrammed in a manner expected for HPV16. In addition, these cells support a variety of life cycle markers including increased proliferation, enhanced incorporation of BrdU, enhanced DNA damage, E1^E4 expression and amplification of the viral genome in the upper layers of the differentiated epithelium as demonstrated by fluorescent in situ hybridization (FISH). Having the parental N/Tert-1 cells allows comparison of interaction of genes with the HPV16 life cycle. To exploit this system we used CRISPR/Cas9 editing to reduce WRN protein expression in N/Tert-1 and N/Tert-1+HPV16 cells. The parental N/Tert-1 cells demonstrated an enhanced proliferative phenotype in the absence of WRN, with increased DNA damage. In N/Tert-1+HPV16 cells lacking WRN there was an increased basal cell layer proliferation phenotype with enhanced DNA damage throughout the epithelium. There was also enhanced numbers of cells supporting viral genome amplification and an overall increase in viral DNA replication as determined by RT-qPCR. Overall the results demonstrate that WRN expression is required for normal epithelial cell differentiation, and also for controlling the HPV16 life cycle in differentiating keratinocytes. The results demonstrate that WRN is a restriction factor for HPV16, controlling the DNA damage induced by the virus and the overall replication levels of the virus.

## Results

### An inverse correlation between SIRT1 and WRN expression in HPV16 positive cervical lesions

Our recent work demonstrated that deacetylation of WRN by SIRT1 promotes binding of WRN to E1-E2 replicating DNA (25). Removal of SIRT1 resulted in hyper-acetylation and stabilization of WRN. In addition, removal of WRN expression from C33a cells by CRISPR/Cas9 resulted in a similar phenotype to that of SIRT1 deletion; increased DNA replication that has a reduced fidelity (25). Therefore, a SIRT1 controls the recruitment of WRN to E1-E2 replicating DNA, and depletion of SIRT1 or WRN results in mutagenic HPV16 E1-E2 DNA replication. These data suggest that manipulation of the SIRT1-WRN axis by HPV may promote cervical disease progression.

To investigate this, we utilized cervical liquid-based cytology samples from a cohort of HPV16+ patients representing the progression of cervical disease (CIN1-CIN3) and compared this to HPV-normal cervical tissue for SIRT1/WRN protein and mRNA expression (50, 51). Figure 1A demonstrates that decreased *WRN* mRNA expression negatively correlated with cervical disease progression through CIN1-CIN3; in contrast, *SIRT1* expression positively correlated with cervical disease progression. In agreement with these data, WRN protein expression negatively correlated with cervical disease expression, whereas SIRT1 protein expression positively correlated with cervical disease progression (Figure 1C; quantification of a larger subset in Figure 1C). Our previous data demonstrated that depletion of SIRT1 stabilized WRN protein by promoting its acetylation (25); when we compared the protein expression of SIRT1 and WRN in matched samples, we observed that WRN expression negatively corelated with SIRT1 expression during cervical disease progression (Figure 1D; CIN1, r = -0.6843, p = 0.005; CIN2, r = -0.6662, p = 0.007; CIN3, r = -0.5200, p = 0.047). Importantly, this was not observed at the mRNA level (Figure 1E). Therefore, although there is an overall trend for WRN mRNA downregulation and SIRT1 mRNA upregulation in the CIN samples (Figure 1A), this was not directly correlative in the individual samples. Together, these data support our previous studies demonstrating that SIRT1 modulates WRN expression in a post-translational manner (i.e. by de-acetylation), and suggest that the SIRT1-WRN axis may contribute to cervical disease progression.

**Figure 1.**
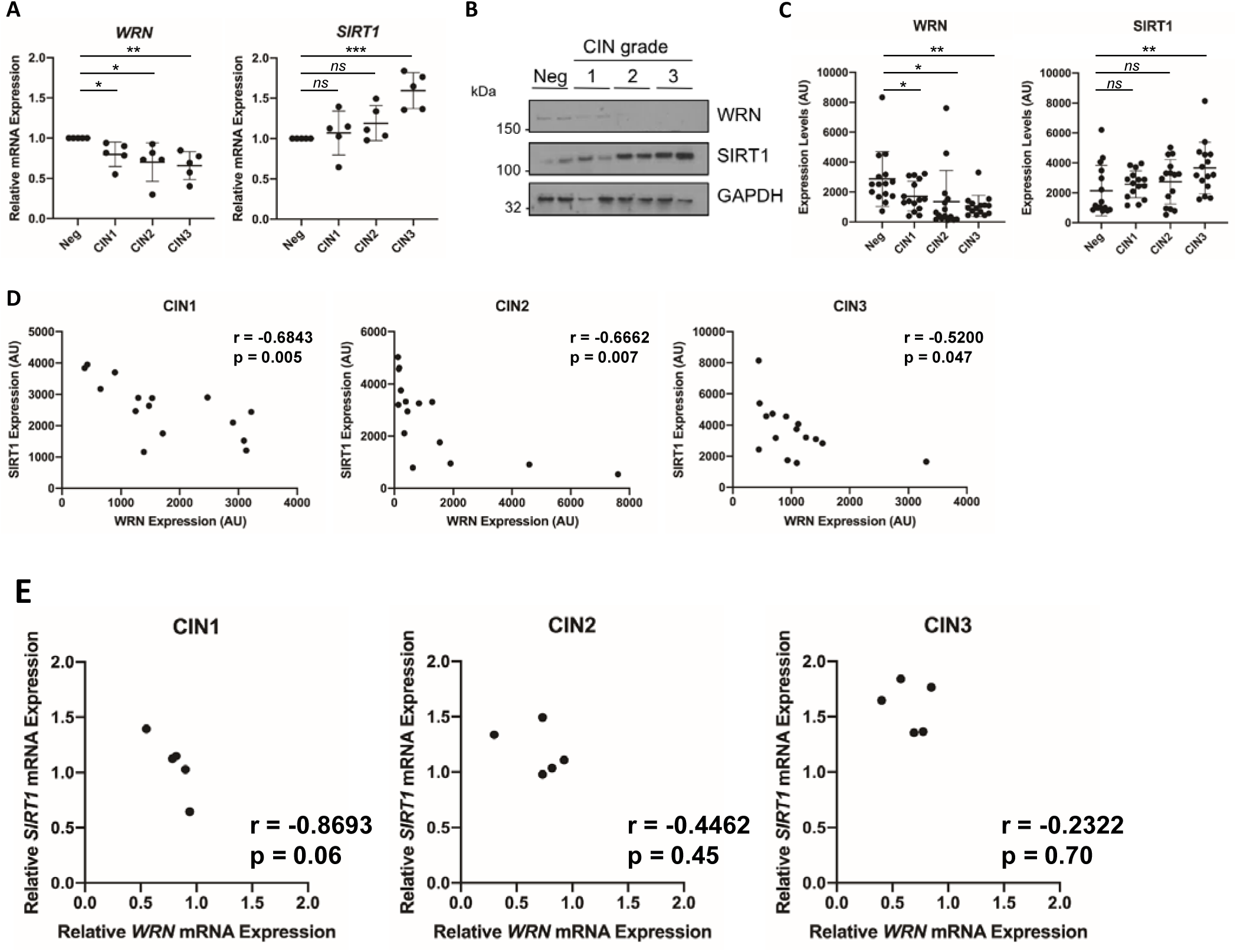
WRN and SIRT1 expression are differentially regulated in cervical disease and negatively correlated at the protein level. **A)** Scatter dot plot of RT-qPCR analysis *WRN* and *SIRT1* mRNA expression from a panel of cervical cytology samples representing CIN legions of increasing grade. Five samples from each clinical grade (neg, CIN I–III) were analyzed and mRNA expression was normalized to the negative samples. Expression was normalized against *U6* mRNA expression. **B)** Representative western blots from cytology samples of CIN lesions of increasing grade analyzed for WRN and SIRT1 expression. GAPDH served as a loading control. **C)** Scatter dot plot of densitometry analysis of a panel of cytology samples for WRN and SIRT1 expression. Fifteen samples from each clinical grade (neg, CIN I–III) were analyzed by western blot and densitometry analysis was performed using ImageJ. **D)** Scatter dot plot analysis of WRN and SIRT1 expression in matched cervical cytology samples in CIN1-3. **E)** Scatter dot plot analysis of *WRN* and *SIRT1* mRNA expression in matched cervical cytology samples in CIN1-3. Correlation between WRN and SIRT1 expression was calculated using Pearson correlation coefficient (r) analysis. Error bars represent the mean ± standard deviation. ns-not significant, * p < 0.05, * p < 0.01, *** p < 0.001 (Student’s *t*-test).

### Generation of WRN depleted N/Tert-1 and N/Tert-1+HPV16 cells using CRISPR/Cas9

To determine the role of WRN in the HPV16 life cycle, N/Tert-1 and N/Tert-1+ HPV16 WRN knockout cells were generated using CRISPR/Cas9 targeting (Figure 2A). Figure 2A demonstrates that the CRISPR/Cas9 targeting of WRN reduced the expression of WRN protein in a heterogeneous cellular pool. To further confirm WRN depletion, we used MTT assay using a WRN helicase inhibitor, the rationale being that the WRN depleted cells should have enhanced resistance to the drug as the cells will be reprogrammed to grow in the absence of WRN. Figure 2B, left panel, demonstrates that this is the case for the N/Tert-1 cells; the WRN depleted cells (N/Tert-1-WRN) have a statistically significantly enhanced resistance to NSC19630. In N/Tert-1+HPV16 cells (right panel), there was little difference between the N/Tert-1+HPV16 and N/Tert-1+HPV16-WRN in their response to NSC19630. This suggests that the WRN present in the N/Tert-1+HPV16 cells is attenuated in function; the IC50 for N/Tert-1+HPV16 and N/Tert-1+HPV16-WRN is similar to N/Tert-1-WRN. This is similar to other DNA damage response proteins that are manipulated by E6 and E7 to abrogate their wild type function and recruit them to the viral DNA to promote viral DNA replication (52). This disruption of the wild type function of these DNA damage response proteins also blocks their ability to signal to the host DNA that there is an active DDR, which would ordinarily promote a cell cycle arrest. The N/Tert-1+HPV16 cells have an active DDR turned on (49).

**Figure 2.**
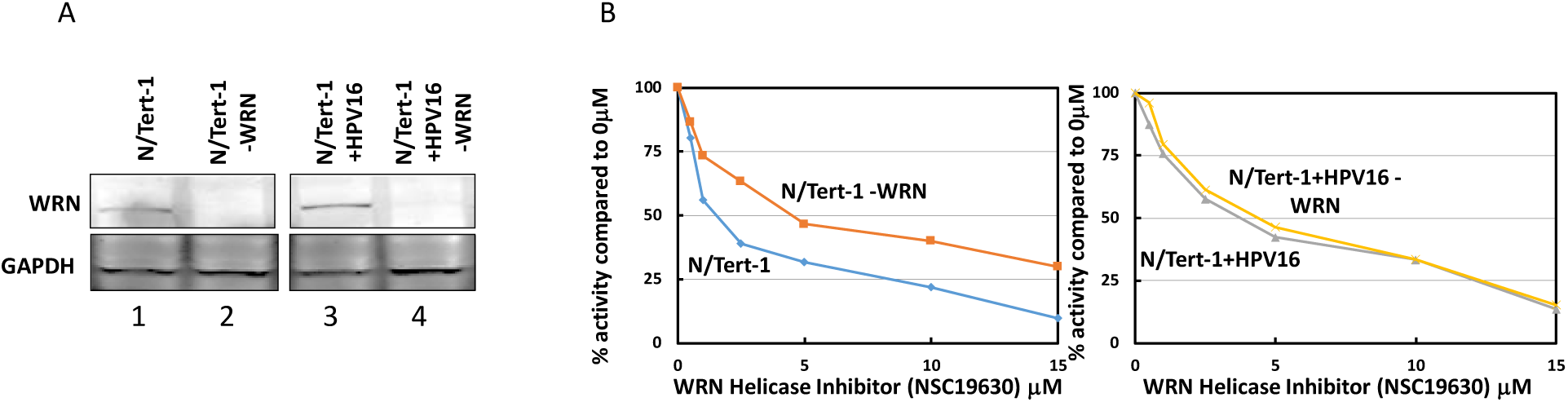
Depletion of WRN from N/Tert-1 and N/Tert-1+HPV16 cells using CRISPR/Cas9. **A)** A plasmid targeting WRN using CRISPR/Cas9 was transfected into N/Tert-1 and N/Tert-1+HPV16 cells and pooled cell lines selected following puromycin treatment. We have used this targeting plasmid previously (25). Western blotting revealed reduced WRN expression in the targeted cell lines (compare lanes 2 and 4 with 1 and 3). **B)** The indicated cell lines were treated with the WRN helicase inhibitor NSC19630 and cell survival estimated using an MTT assay. With the N/Tert-1 and N/Tert-1-WRN cells there as a statistically significant difference in the response to the drug from the 1μM concentration forward (p-value <0.05); for the N/Tert-1+HPV16 pair there was no statistically significant difference between N/Tert-1+HPV16 and N/Tert-1+HPV16-WRN. This was determined using a student t-test.

### WRN depletion enhances the proliferation of N/Tert-1 cells

In order to investigate the role of WRN in regulation of cell proliferation and differentiation of epithelial cells, and whether WRN depletion altered the HPV16 life cycle during this process, N//Tert-1, N/Tert-1-WRN, N/Tert-1+HPV16 and N/Tert-1+HPV16-WRN were subjected to organotypic raft culture experiments. Figure 3A presents images from H&E staining of the cultures and are representative of two independent raft experiments for each cell line. The presence of HPV16 enhances the thickness of the N/Tert-1 cells as expected (compare N/Tert-1 with N/Tert-1+HPV16; independent rafts were scanned using a Vectra Polaris machine that calculated their overall thickness (“height”)). Furthermore, the depletion of WRN had a marked effect on the raft culture, increasing the thickness of the raft substantially (Figure 3B). This demonstrated the significance of the enhanced thickening of the epithelium following WRN depletion or expression of HPV16.

**Figure 3.**
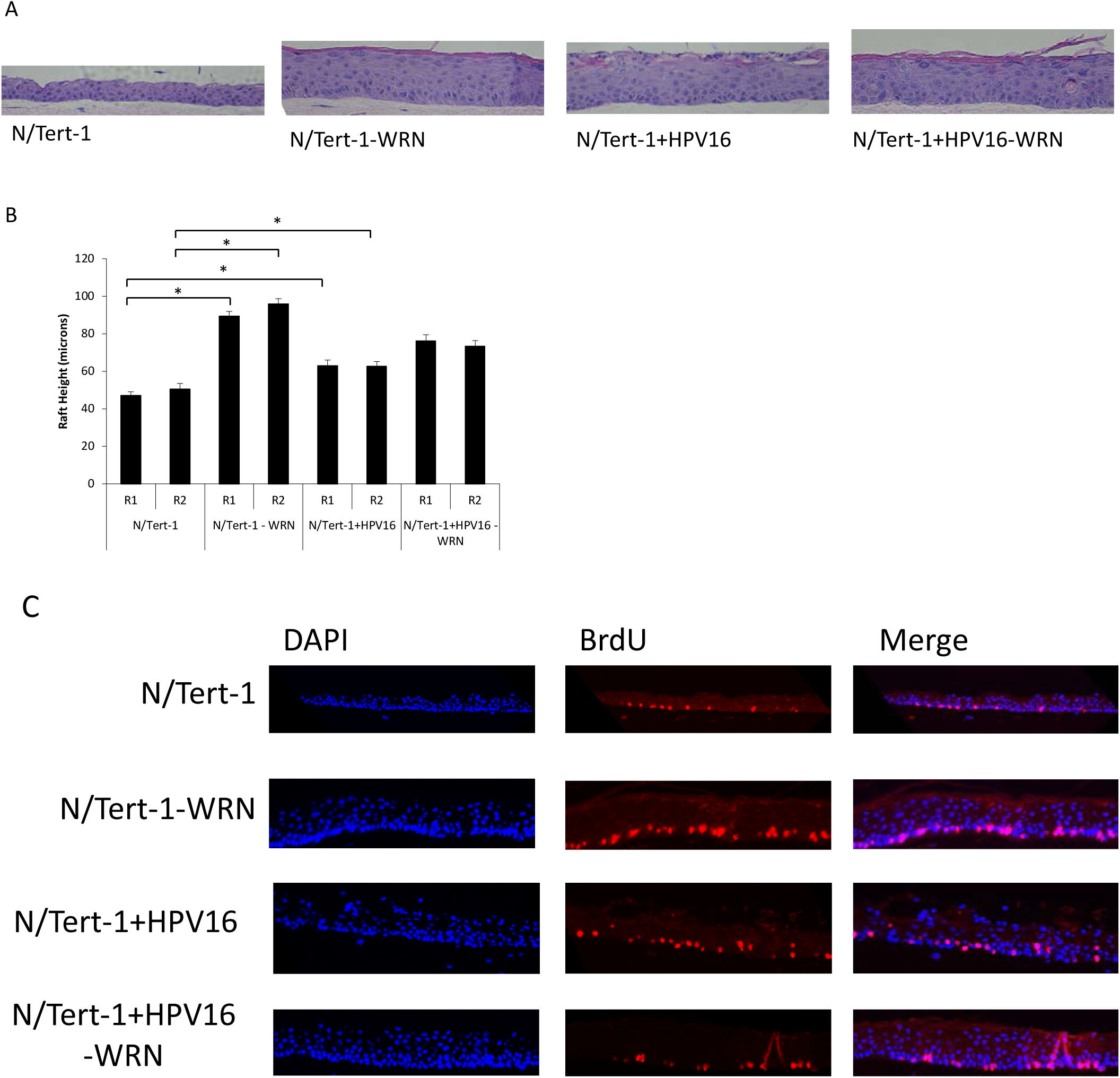
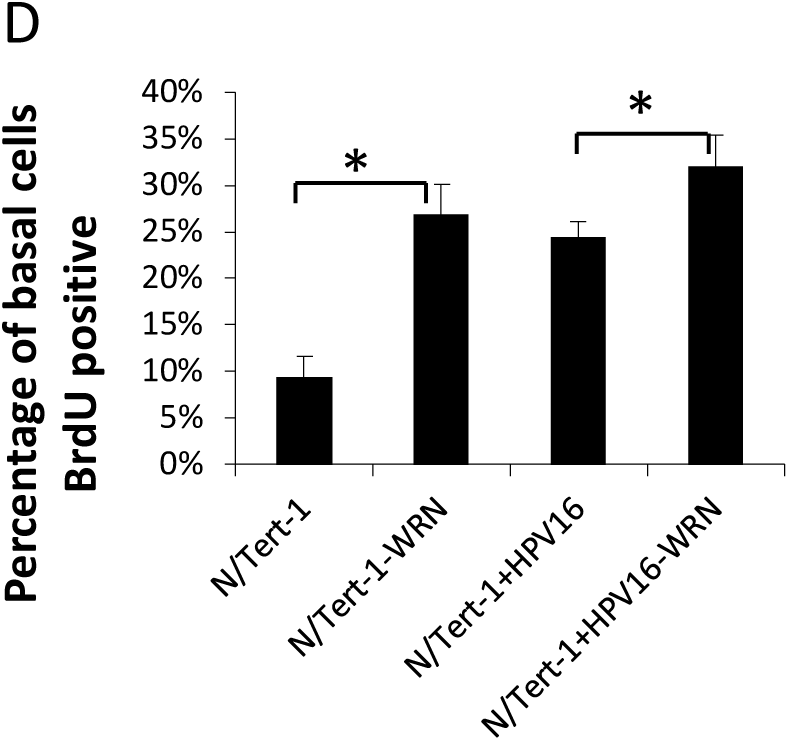
Knockdown of WRN increases proliferation during differentiation. **A)** WRN CRISPR and control cell lines (Figure 2) were differentiated for 13 days by organotypic raft culture. Fixed 4μM sections were subject to H&E staining and representative samples are shown. **B)** Two H&E stained sections from two individual rafts were imaged and measurements were taken at 100 micron intervals across the length of each section. When compared to parental lines, N/Tert-1 WRN CRISPR lines were hyperproliferative. HPV16 increase proliferation that was not enhanced significantly following WRN removal. **C)** Nucleoside analog BrdU was included in media for the final 16 hours of organotypic raft culture. Fixed sections were stained for BrdU incorporation. **D)** The number of BrdU positive cells was counted using a Vectra^®^ Polaris™ Imaging System; whole stained sections were scanned computationally and the intensity calculated compared to a negative background control (secondary antibody only) and a positive localisation control (DAPI). The same imaging parameters were used for each slide. Three sections from two individual rafts were subject to analysis. BrdU incorporation and quantification revealed that proliferation of basal cells is enhanced in both N/Tert-1 and N/Tert-1+HPV16 cells in the absence of WRN. Error bars indicate standard error of the mean and * indicates p<0.05.

To investigate the effects of WRN depletion on proliferation, the raft cultures were labelled with BrdU for 16 hours prior to harvesting. Staining with a BrdU antibody was then carried out (Figure 3C) and the results quantitated (Figure 3D); the images are representative and the quantitation an average from the independent raft experiments. N/Tert-1-WRN cells have an enhanced basal layer proliferative capacity as they have incorporated more BrdU than the N/Tert-1 cells. Similarly, HPV16 enhances BrdU incorporation in the N/Tert-1 cells, as we previously demonstrated (48). While not clearly evident on the images, the lack of WRN also increased BrdU incorporation into the N/Tert-1+HPV16 cells. Although difficult to quantitate, there was an increase of suprabasal cells staining positive for BrdU in the N/Tert-1-WRN, N/Tert-1+HPV16 and N/Tert-1+HPV16-WRN when compared with N/Tert-1. Overall, these results demonstrate that the absence of WRN increases cell proliferation in N/Tert-1 cells and that basal cell proliferation is enhanced in both N/Tert-1 and N/Tert-1+HPV16 cells in the absence of WRN.

We next investigated differentiation markers in the cells to determine whether WRN depletion was having an effect on this process by staining for Keratin 10 and Involucrin (Figure 4). In the N/Tert-1-WRN and N/Tert-1+HPV16 cells there is a slight change in Involucrin staining as basal and immediately supra-basal cells have a delayed expression compared with N/Tert-1. This indicates a short delay in differentiation, but that the overall differentiation process is occurring in the N/Tert-1-WRN and N/Tert-1+HPV16 cells. Therefore, the enhanced proliferation is not due to a failure of the cells to differentiate.

**Figure 4.**
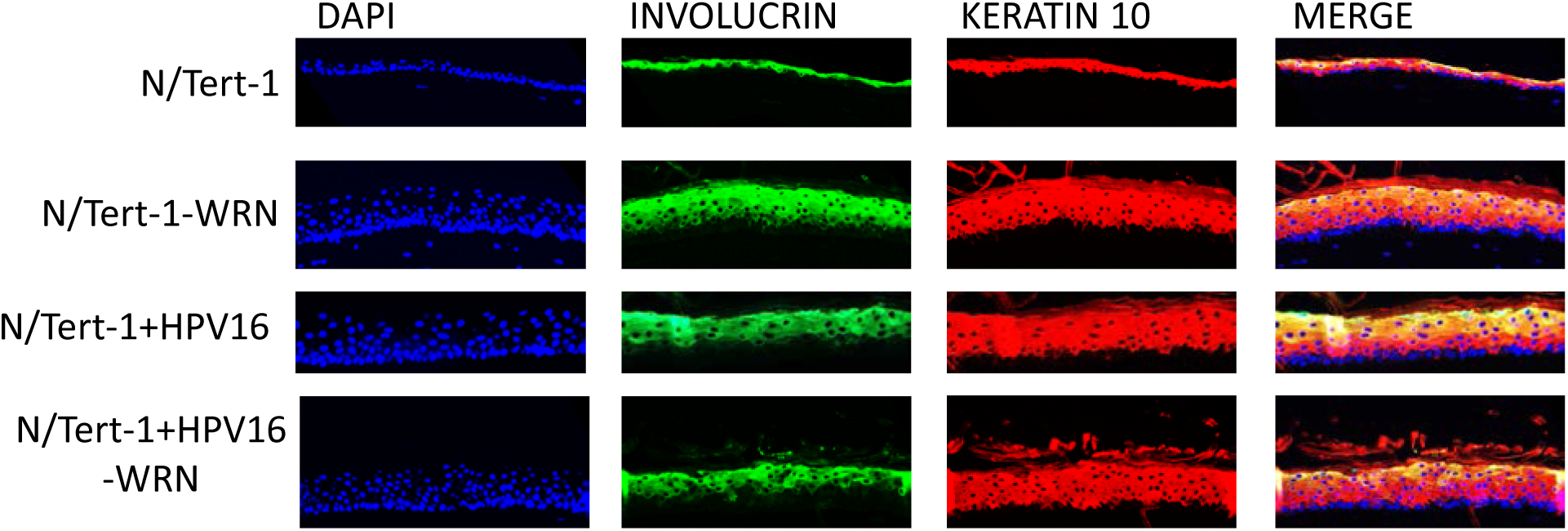
Knockdown of WRN has no affect upon differentiation of N/Tert-1 cell lines. Sections were stained with the indicated antibodies. Both proteins are markers of terminal differentiation, and are observed in suprabasal cells, and absent from basal cells, as expected during normal organotypic raft culture. DNA was stained with 4’,6-diamidino-2-phenylindole (DAPI). Images are representative of two individual rafts grown per cell line.

### WRN depletion alters the HPV16 life cycle in N/Tert-1 cells

We next looked at a variety of life cycle markers in the rafts to determine whether the depletion of WRN interferes with the HPV16 life cycle in N/Tert-1 cells. γH2AX staining has been used as a surrogate marker for HPV16 replication during the viral life cycle, particularly during the amplification stage of the life cycle in the upper layers of the epithelium (21). We stained our N/Tert-1 panel of cells with γH2AX and Figure 5A gives representative images of the results. There was a residual level of γH2AX staining in N/tert-1 cells; however, WRN depletion increases the number of cells that are γH2AX positive. This demonstrates that an increase in DNA damage accompanies the increased proliferation induced by the removal of WRN (Figure 3). As expected, the presence of HPV16 induced increased γH2AX staining in the upper layers of the differentiating epithelium where genome amplification occurs, with weaker but consistent staining in the basal layers of the epithelium. Depletion of WRN in the N/Tert-1+HPV16 cells increases the number of cells staining positive for γH2AX; in particular, there is an increase in basal layer cells staining positive, with corresponding increase in the intensity of the signal. We quantitated the γH2AX positive cells using the Vectra Polaris machine and Figure 5B demonstrates that depletion of WRN or the presence of HPV16 increases the overall γH2AX staining percentage. Staining of the basal layers demonstrates a large increase in γH2AX staining in the absence of WRN, and this is also observed in the presence of HPV16 (Figure 5C). Overall, the results demonstrate that WRN depletion results in elevated γH2AX staining in the absence and presence of HPV16 in N/Tert-1 cells.

**Figure 5.**
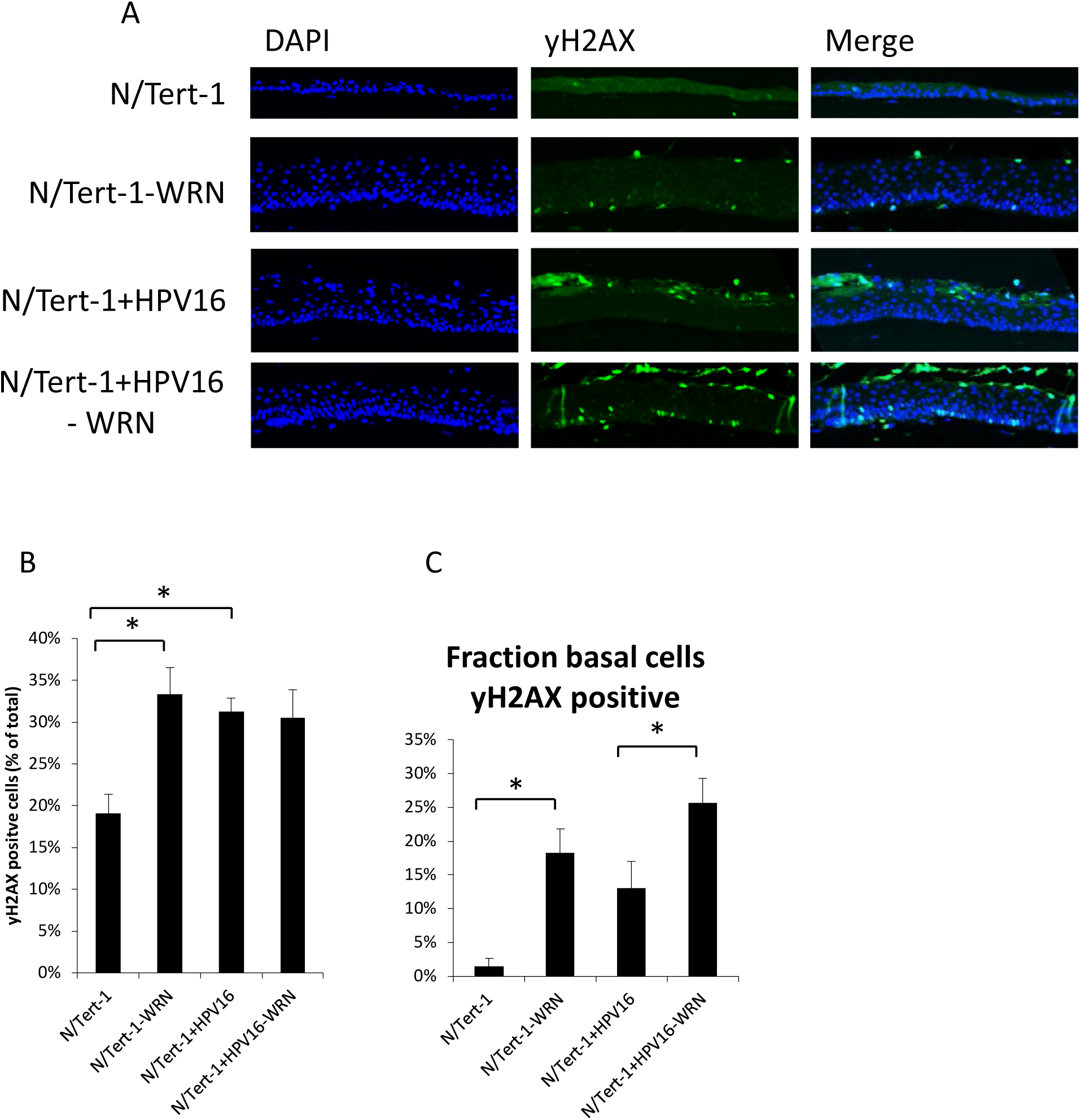
Knockdown of WRN increases DNA damage. **A)** To visualize DNA damage throughout differentiated sections, staining for γH2AX was carried out; phosphorylation of H2AX at Ser-139 (γ-H2AX) correlates with sites of DNA damage. Cellular DNA was stained with 4’,6-diamidino-2-phenylindole (DAPI). **B)** The number of yH2AX positive cells was measured using a Vectra^®^ Polaris™ Imaging System; whole stained sections were scanned computationally and the intensity calculated compared to a negative background control (secondary antibody only) and a positive localisation control (DAPI). **C)** The number of yH2AX positive cells was calculated as a fraction of the number of basal cells, where only the basal yH2AX counts were included in quantification. Three sections from two individual rafts were subject to analysis. Error bars indicate standard error of the mean and * indicates p<0.05.

Using fluorescent in situ hybridization (FISH) to detect the HPV16 genome, we investigated the regulation of the viral genome during differentiation of the N/Tert-1 cells. Figure 6A demonstrates the usual FISH signal detected in the upper layers of the differentiated epithelium, representative of amplification of the viral genome. WRN depletion resulted in detection of the viral genome throughout the differentiating epithelium with a concentration in the upper layers remaining. Using our Vectra Polaris machine, we quantitated the staining for the number of cells that are HPV16 positive and the intensity of the staining (Figure 6B).

**Figure 6.**
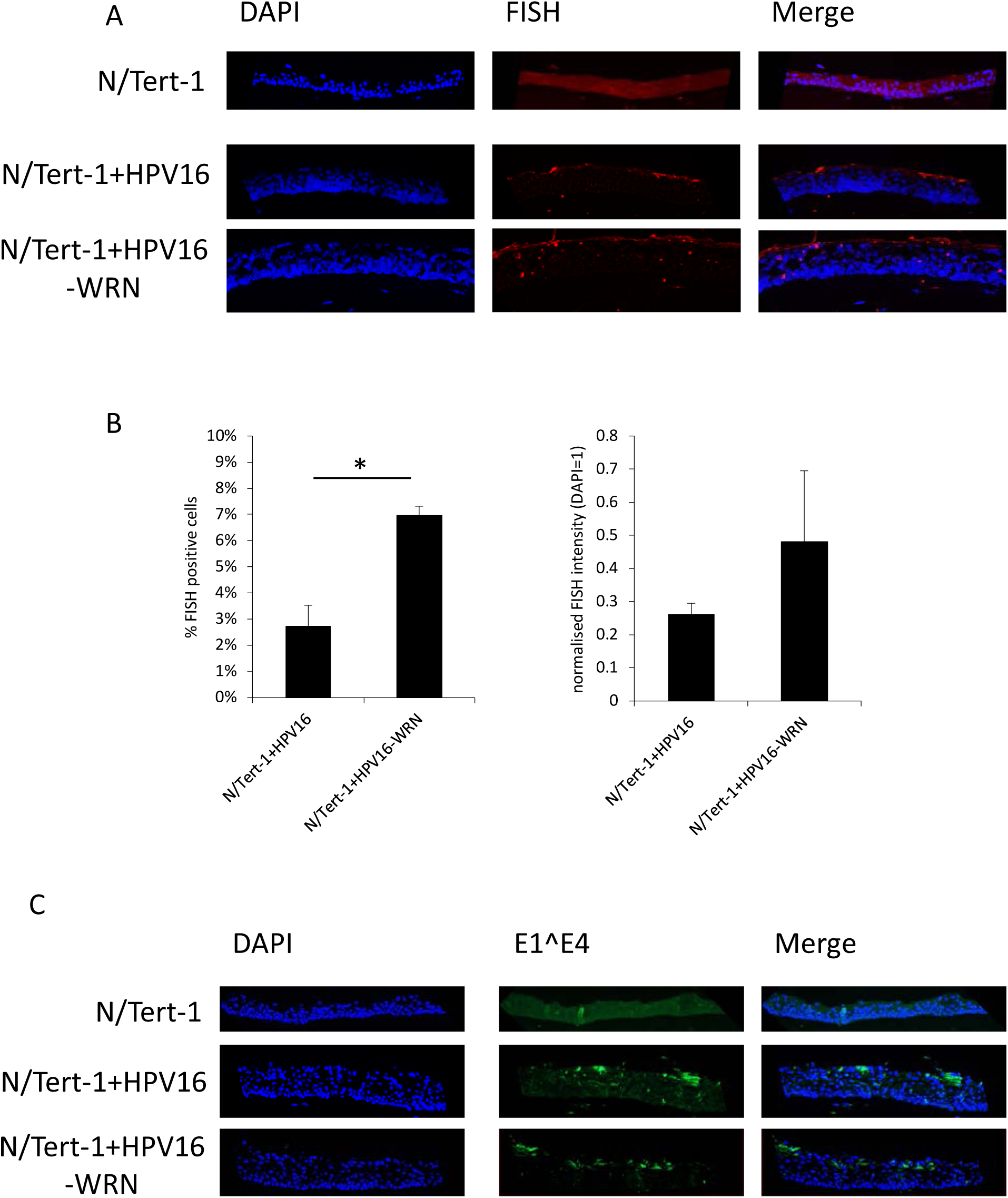
Depletion of WRN increases the number of cells supporting viral replication in the differentiating epithelium. **A)** N/Tert-1+HPV16 and N/Tert-1+HPV16 CRISPR WRN cells were differentiated in culture, formalin fixed, paraffin embedded, sectioned and then stained for HPV16 genomes using DNA-FISH. Images shown are representative images of HPV-FISH in differentiated culture. **B)** Both the percentage of HPV16 positive cells and the intensity of fluorescence was quantified (B) using Vectra^®^ Polaris™ Imaging System. The intensity of fluorescence was calculated compared to a negative background control (N/Tert-1) and the same imaging parameters were used for each slide. Three sections from two individual rafts were subject to analysis, error bars indicate standard error of the mean and * indicates p<0.05. **C)** Sections were stained with HPV16 E1^E4 to investigate whether a productive viral life was occuring; the E1^E4 spliced variant is expressed during the late stages and upper layers of epithelium. Here, we observe that E1^E4 expression occurs regardless of WRN depletion.

We next investigate expression of E1^E4 staining in the epithelium to determine whether this late stage marker of the viral life cycle is disrupted following depletion of WRN. Figure 6C demonstrates that this was not the case, the depletion of WRN had no significant effect on the expression of E1^E4. The E1^E4 staining is similar to that observed by others (53).

The expression of E1^E4 demonstrates that episomal genomes remain in the N/Tert-1+HPV16-WRN cells. We carried out Southern blots and real-time PCR to investigate further the status of the viral genomes in the cells. A single cutter on the viral genome (Sph1) generated an 8kbp band in all samples irrespective of monolayer, raft or WRN status (Figure 7A). Treatment with HindIII, a non-cutter of the viral genome, resulted in a uniform detection of nicked DNA (Figure 7B). We did not observe supercoiled DNA on this blot, perhaps due to the DNA preparation protocol. The uniform nature of the bands in Figures 7A and B, combined with the E1^E4 status, demonstrates a large presence of episomal viral genomes in the presence and absence of WRN. As Southern blots are at best semi-quantitative, we carried out real-time PCR on the DNA harvested from the rafts. We monitored for the levels of E6 DNA, and Figure 7C demonstrates that there is a small but significant increase in viral genomes following WRN depletion. This is in agreement with the FISH data (Figure 6). These results demonstrate that the HPV16 genome remains episomal following WRN depletion and that there is an increase in viral genome copy number.

**Figure 7.**
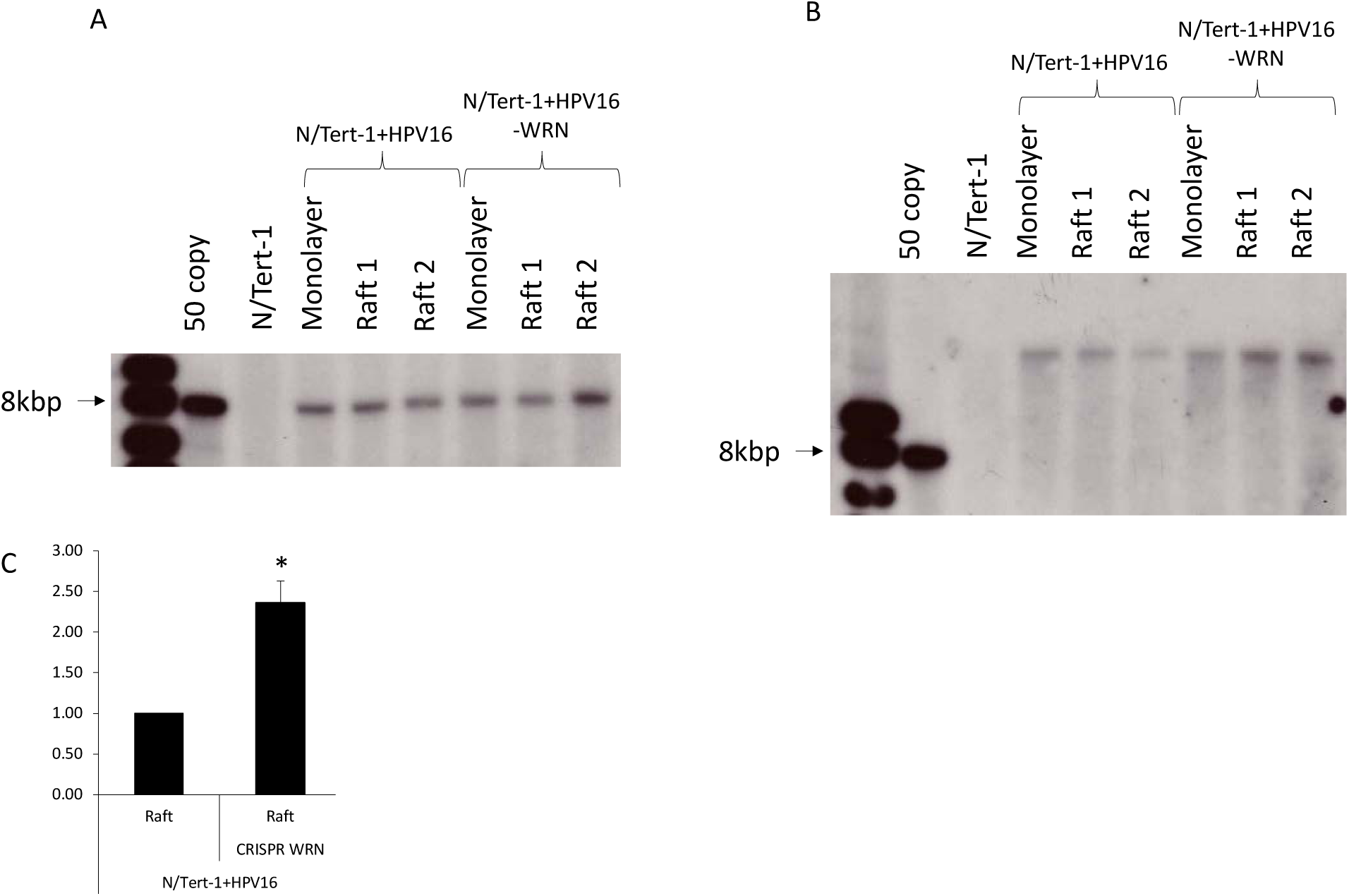
WRN depletion does not dramatically alter the status of the HPV16 genome. **A&B)** DNA was extracted from monolayer and raft cultures, parental and CRISPR WRN cells. This DNA was then digested with a non-cutter (*Sph*I, **A**) or a single cutter (*Hind*III, **B**) of the HPV16 genome, and subject to Southern blot analysis using an HPV16 genome probe. **C)** The same DNA was subject to PCR analysis using E6 primers. To calculate HPV amplification, E6 ΔCt values were first calculated relative to mitochondrial housekeeping gene and ΔΔCt relative to the parental cells line (N/Tert-1+HPV16). Error bars indicate standard error of the mean of two experimental repeats and * indicates p<0.05.

## Discussion

Our previous work demonstrated that WRN was a substrate for SIRT1, in agreement with other reports (25, 44, 45). The consequence of SIRT1 depletion by CRISPR/Cas9 was an enhanced acetylation and stabilization of WRN; however, acetylated WRN is unable to complex with damaged and E1-E2 replicating DNA, perhaps due to the negative charge associated with acetylation (25). In the absence of SIRT1, there was a reduction of HPV16 E1-E2 replication fidelity and a decreased recruitment of WRN to the replicating DNA. The role of WRN in regulation of HR could at least partially account for the mutagenic replication in the absence of SIRT1. Indeed, following WRN depletion by CRISPR/Cas9 targeting there was also a reduction in the fidelity of E1-E2 DNA replication. Our mechanistic studies therefore suggested a SIRT1-WRN axis controlling viral DNA replication fidelity. There is also an important role for SIRT1 during the HPV31 life cycle; removal of SIRT1 blocked viral genome amplification (30). This study demonstrated alteration in viral genome histone modifications that could have contributed to the failure to amplify the viral genome during epithelial cell differentiation. To investigate the SIRT1-WRN axis further we studied the expression levels of these proteins in naturally occurring cervical HPV positive CIN1, 2 and 3 lesions compared with normal cervical tissue (Figure 1). While there is a tendency towards increasing SIRT1 mRNA levels and decreasing WRN mRNA levels as disease progresses, it is clear that there is an inverse correlation between the SIRT1 and WRN proteins during disease progression. This was not accounted for by mRNA levels as, with individual lesions, there was no clear inverse correlation between expression of SIRT1 and WRN mRNA. Overall, this study suggested that elevated SIRT1 results in decreased WRN acetylation and stability during HPV16 disease progression. The deacetylated WRN, although reduced in stability and therefore detectable levels in these lesions, is much more active at complexing with replicating DNA and promoting HR repair.

To investigate the role of WRN during the viral life cycle, we exploited our N/Tert-1 model. N/Tert-1 are hTERT immortalized human foreskin keratinocytes and we generated cell lines containing HPV16, N/Tert-1+HPV16 (47-49, 54). This system has the direct advantage of being able to manipulate control cells (N/Tert-1) without the viral genome, but with the same genetic lesions as the N/Tert-1+HPV16 cells. Without this comparison, it would be more difficult to determine whether the changes observed in the presence of HPV16 with the genetic lesion is due to the presence of the virus, or due to the effect of the lesion on the epithelium. CRISPR/Cas9 targeting generated N/Tert1-WRN and N/Tert-1+HPV16-WRN; the plasmid used to target WRN has been described previously (25), Figure 2. We tested the effect of a WRN helicase inhibitor, NSC19630, on the viability of wild type and WRN depleted N/Tert-1 and N/Tert-1+HPV16 and observed that the N/Tert-1-WRN cells were more resistant to this drug than the control N/Tert-1, as would be expected. This drug obviously has toxicity and off-target effects as the N/Tert-1-WRN cells were still susceptible to growth attenuation when treated with NSC 19630. Interestingly, in N/Tert-1+HPV16 the absence of WRN made no difference to their growth in the presence of NSC19630. This suggests that, in N/Tert-1+HPV16 cells, the WRN that is present is not functioning correctly. Indeed, the IC50 of the N/Tert-1+HPV16 and N/Tert-1+HPV16-WRN are both similar to N/Tert-1-WRN. This suggests that, in HPV16 positive cells, WRN function on the host genome is attenuated, and that it is recruited to the viral genome to assist with viral replication. This is similar to other DNA repair factors; E6 and E7 manipulate these factors so that they do not interact correctly with the host genome (52). Using this mechanism, the virus would allow the presence of an active DDR to promote the viral life cycle but prevent the host cell detecting this damage by subverting the ability of host factors to signal the damage to the host replication machinery. This would allow proliferation with an active DDR in HPV16 positive cells. We are currently investigating the mechanism that HPV16 uses to subvert the normal function of WRN in HPV16 positive cells.

To determine the effect of WRN depletion on the viral life cycle, we subjected the cell lines to organotypic raft culture, followed by investigation of markers of the viral life cycle. Figure 3 demonstrates that, in the absence of WRN, there is a thickening of the N/Tert-1 differentiating tissue to a level similar to rafts containing HPV16. There was no further increase in epithelial proliferation in N/Tert-1+HPV16 cells in the absence of WRN. Werner syndrome patients suffer from progeria but are also predisposed to certain cancers, therefore WRN is a tumor suppressor protein (31, 55, 56). The thickening of the N/Tert-1 epithelium in the absence of WRN is a proliferative signal, a hallmark of the absence of a tumor suppressor. To demonstrate an enhanced proliferation in the absence of WRN, we labeled our cultures with BrdU 16 hours before harvesting the rafts, allowing us to stain for BrdU expression to investigate the proliferative status of the epithelium. Figure 3 demonstrates that the removal of WRN enhances BrdU incorporation, in both N/Tert-1 and N/Tert-1+HPV16 cells. The alteration in cell proliferation was not due to an obvious attenuation of cell differentiation, as expression of the differentiation markers Involucrin and Keratin 10 were not altered in the absence of WRN (Figure 4).

A hallmark of HPV16 infection is activation of the DDR, detected in differentiated epithelium; expression of γH2AX can be used to detect the induced damage (21). Figure 5 demonstrates that there is an increase in γH2AX signal in N/Tert-1 and N/Tert-1+HPV16 following WRN depletion, demonstrating that the increased proliferation induced by WRN depletion is accompanied by elevated DNA damage. Such a phenotype would promote cell transformation, perhaps an expected phenotype following removal of expression of a tumor suppressor protein.

Finally, we moved on to investigate whether depletion of WRN influences the levels and status of the viral genome in N/Tert-1+HPV16 cells (Figures 6 and 7). Using FISH, we demonstrated an increase in the number of HPV16 positive cells with accompanying increase in fluorescent intensity (Figure 6A&B). A clear phenotype of WRN depletion was also an increase in the number of FISH signals observed in the mid-layer of the differentiating epithelium. In Figure 6A, N/Tert-1+HPV16 fluorescent staining is concentrated in the upper layers of the epithelium where viral genome amplification occurs, as we have reported previously (49). However, in N/Tert-1+HPV16-WRN, there are a significant number of fluorescent cells in the mid-layer of the epithelium. Another marker of late stages of the HPV16 life cycle is expression of E1^E4 towards the upper layer of the epithelium, not disrupted by the lack of WRN expression (Figure 6C). This suggested that the viral DNA remains episomal in the absence of WRN, and Southern blotting demonstrated this to be the case, both in monolayer and in the differentiating epithelium (Figure 7). While these blots do not demonstrate the lack of integration (which would be a random event in a pool and therefore would not show up as an individual event/band on a Southern) in the absence of WRN, they do demonstrate that there remains a significant level of episomal viral genomes. Southern blots are semi-quantitative, but real-time qPCR demonstrated an increase in viral genomes in the absence of WRN, agreeing with the FISH data from Figure 6.

Overall, the results presented here demonstrate that WRN controls the proliferation of differentiating epithelium; the absence of WRN elevates proliferation and DNA damage irrespective of HPV16 status. In the presence of HPV16, the absence of WRN gives an additive effect on cell proliferation and DNA damage. The one phenotype revealed in N/Tert-1+HPV16-WRN is an improper amplification of the viral genome in the mid-layers of the differentiating epithelium resulting in increased levels of viral DNA. Therefore, WRN functions as a restriction factor for HPV16 as it controls replication during the differentiating epithelium. This is similar to SAMHD1, another DNA replication and repair factor involved in homologous recombination (57, 58). Recently, we demonstrated that, although SAMHD1 levels are downregulated by HPV16, full deletion of SAMHD1 using CRISPR/Cas9 results in an increased amplification of the viral genome during differentiation (48). Therefore, there is a delicate interaction between HPV16 and WRN/SAMHD1; these proteins must be downregulated to promote viral replication but are required for optimal replication, their depletion results in enhanced amplification of the viral genome. Whether this amplification is of high fidelity remains to be determined, depletion of SIRT1 and WRN results in low fidelity replication in our HPV16 E1-E2 C33a cell model (25, 26). Both SAMHD1 and WRN can regulate the function of MRE11, a component of the MRN complex (MRE11, RAD50, NBS1) (57-59). MRN function is required for amplification of HPV31 during epithelial differentiation, therefore has the opposite effect to WRN and SAMHD1 where depletion boosts replication (22). We are currently investigating whether the WRN and SAMHD1 phenotypes are mediated via disrupted MRE11 activity.

The SIRT1-WRN axis is a crucial one for the viral life cycle that controls the levels of viral DNA replication. Future studies will concentrate on investigating the role of WRN and partner proteins in the viral life cycle, with the ultimate goal of being able to block viral replication and/or kill the infected cell.

## Materials and Methods

### Cell line, plasmids, and reagents

N/Tert-1 and N/Tert-1+HPV16 cells were grown and maintained in K-SFM media containing 1% (vol/vol) penicillin-streptomycin mixture and 4 µg/ml hygromycin B at 37°C in a 5% CO2 incubator (25). For WRN knockout CRISPR, WRN double-nickase plasmid (h) (sc-401860-NIC) was purchased from Santa Cruz. N/Tert-1 and N/Tert-1+HPV16 WRN–/– pools were generated as described for the SIRT1 knockout cells (26).

### MTT cell proliferation assay

MTT Kit (ATCC 30–1010 K) was purchased from ATCC and was used for the cell proliferation assay as mentioned in Das *et al*., 2017 (52). Briefly the cells were plated in a 96 well tissue culture plate and treated with WRN helicase inhibitor (NSC19630) for a specified time period. After washing MTT reagent was added for the formation of the purple crystals which were later dissolved using a detergent solution. Later absorbance was measured at 570 nm and the data was presented as percentages of the control.

### Cervical cytology samples

Cervical cytology samples were obtained from the Scottish HPV Archive (http://www.shine/mvm.ed.ac.uk/archive.shtml), a biobank of over 20,000 samples designed to facilitate HPV associated research. The East of Scotland Research Ethics Service has given generic approval to the Scottish HPV Archive as a Research Tissue Bank (REC Ref 11/AL/0174) for HPV related research on anonymised archive samples. Samples are available for the present project though application to the Archive Steering Committee (HPV Archive Application Ref 0034). RNA and protein were extracted from the samples using Trizol and analyzed as previously described (50, 51).

### Quantitative reverse transcriptase real-time PCR on cervical cytology RNA samples

Total RNA was extracted using the E.Z.N.A. Total RNA Kit I (Omega Bio-Tek) according to the manufacture’s protocol. One μg of total RNA was DNase treated following the RQ1 RNase-Free DNase protocol (Promega) and then reverse transcribed with a mixture of random primers and oligo(dT) primers using the qScriptTM cDNA SuperMix (Quanta Biosciences) according to instructions. RT-qPCR was performed using the QuantiFast SYBR Green PCR kit (Qiagen). The PCR reaction was conducted on a Corbett Rotor-Gene 6000 (Qiagen) as follows: initial activation step for 10 min at 95°C and a three-step cycle of denaturation (10 sec at 95°C), annealing (15 sec at 60°C) and extension (20 sec at 72°C) which was repeated 40 times and concluded by melting curve analysis. The following primers were used: *WRN* Forward 5’-GCATGTGTTCGGAAGAGTGTTT -3’, *WRN* Reverse 5’-TGACATGGAAGAAACGTGGAA -3’; *SIRT1* Forward 5’-TGCTGGCCTAATAGAGTGGCA -3’, *SIRT1* Reverse 5’-CTCAGCGCCATGGAAAATGT -3’. mRNA expression was normalized against *U6* expression using the following primers: *U6* Forward 5’-CTCGCTTCGGCAGCACA -3’, *U6* Reverse 5’-AACGCTTCACGCATTTGC -3’. The data obtained was analyzed according to the ΔΔCt method using the Rotor-Gene 6000 software (60).

### Western blotting

Total protein was resolved by SDS-PAGE (10–15% Tris-Glycine), transferred onto Hybond nitrocellulose membrane (Amersham biosciences) and probed with antibodies specific for WRN (D-6; sc-376182, Santa Cruz Biotechnology (SCBT)), SIRT1 (B-7; sc-74465, SCBT) and GAPDH (G-9; sc-365062, SCBT). Western blots were visualized with species-specific HRP conjugated secondary antibodies (Sigma) and ECL (Thermo/Pierce). Densitometry analysis was performed using ImageJ analysis software (NIH, USA).

### Organotypic raft culture

N/Tert-1 and N/Tert-1+HPV16 cells were differentiated via organotypic raft culture as described previously (refs). Briefly, cells were seeded onto type 1 collagen matrices containing J2 3T3 fibroblast feeder cells. Cells were then grown to confluency atop the collagen matrices, which were then lifted onto wire grids and cultured in cell culture dishes at the air-liquid interface, with media replacement on alternate days. 16 hours before fixation, media was replaced with fresh media supplemented with 20μM BrdU (final concentration). Following 13 days of culture, rafted samples were fixed with formaldehyde (4% v/v) and embedded in paraffin blocks. Multiple 4μm sections were cut from each sample. For FISH staining, 6μm sections were cut. Sections were stained with hematoxylin and eosin (H&E) and others prepared for immunofluorescent staining as described previously. Fixing and embedding services in support of the research project were generated by the VCU Massey Cancer Center Cancer Mouse Model Shared Resource, supported, in part, with funding from NIH-NCI Cancer Center Support Grant P30 CA016059.

### Immunofluorescence

Antibodies used and relevant dilutions are as follows: Involucrin (1/1000, Santa Cruz Biotechnology), Keratin-10 (1/2000, Santa Cruz Biotechnology), BrdU (1/200, Cell Signaling Technology), phospho-Histone H2AX (Ser139) (1/500, Cell Signaling Technology). Immune complexes were visualized using Alexa 488- or Alexa 595-labeled anti-species specific antibody conjugates (Molecular Probes). Cellular DNA was stained with 4’,6-diamidino-2-phenylindole (DAPI, Santa Cruz sc-3598). Fluorescent in situ hybridization (FISH) staining for HPV16 genomes was performed using DIG-labeled HPV16 genomes, as described previously (references). Microscopy and subsequent quantification performed at the VCU Microscopy Facility, supported, in part, by funding from NIH-NCI cancer center grant P30 CA16059. Image analysis (% Staining and staining intensity) was performed using a Vectra^®^ Polaris™ automated imaging system, whereby whole stained sections were scanned computationally and the intensity calculated compared to a negative background control (secondary antibody only) and a positive localization control (DAPI). The intensity of IF microscopy is a measure of the photons detected from one emission channel (Waters 2009). Intensity was calculated based on the number of photons at a specific location, thus determining the local concentration of fluorophores (secondary antibodies). In this way, this is equivalent to measuring densitometry to estimate protein concentration from western blot. The same imaging parameters were used for each slide and for each sample and two sections from three individually-grown rafts were scanned to generate average values. Immunofluorescence was observed using a LSM 710 Laser Scanning Microscope and ZEN 2011 software (Carl Zeiss).

### Southern Blot

DNA was isolated from monolayer and raft cultures by incubation in HIRT buffer (0.6% SDS, 10 mM EDTA pH 7.5 5 M NaCl) and phenol chloroform extraction, as previously described (Evans *et al.* 2016), and 5 micrograms digested with either *Sph*I or *Hind*III, to linearise the HPV16 genome or leave episomes intact, respectively. All digests included *Dpn*I to ensure that all input DNA was digested and not represented as replicating viral DNA. Digested DNA was separated by electrophoresis of a 0.8% agarose gel, transferred to a nitrocellulose membrane and probed with radiolabeled (32-P) HPV16 genome. This was then visualized by exposure to film for 24 or 72 hours.

### Quantitative PCR on HPV16 DNA samples

DNA was isolated as above and subject to PCR utilizing the SYBR Green Master Mix and 7500 Fast Real-Time PCR System described above. As HIRT buffer is optimized for the isolation of small DNA, mitochondrial DNA was detected as the internal control; F 5’- caggagtaggagagagggaggtaag-3’ R 5’- tacccatcataatcggaggctttgg-3’. HPV16 primers were as follows: E6 F 5’-gagaactgcaatgtttcaggacc-3’ R 5’-tgtatagttgtttgcagctctgtgc-3’; E2 5’-atgcgggtggtcaggtaata-3’ 5’-tcgctggatagtcgtctgtg-3’.

### Statistics

Standard error was calculated from three independent experiments and significance determined using a student’s t-test. For cervical cytology sample analysis, individual samples were plotted with the error bars representing the standard deviation. Significance was determined by two-tailed, unpaired Student’s T-Test. Correlation of WRN and SIRT1 expression was calculated using Pearson correlation coefficient (r) analysis using GraphPad Prism software as previously described (61).

## Acknowledgements

This work was supported by VCU Philips Institute for Oral Health Research and the National Cancer Institute Designated Massey Cancer Center grant P30 CA016059 (IMM). The Medical Research Council (MRC) funded AM (MR/K012665 and MR/S001697/1). ELM received support from the Wellcome Institutional Strategic Support Fund (ISSF) (204825/Z/16/Z). We thank the Scottish HPV Investigators Network (SHINE) for providing HPV positive cytology samples. The funders had no role in study design, data collection and analysis, decision to publish, or preparation of the manuscript.

